# How much information is there for inferring species trees?

**DOI:** 10.64898/2026.04.01.715836

**Authors:** Analisa Milkey, Jessica Chen, Paul O. Lewis

## Abstract

As modern phylogenomics datasets become increasingly large, it is useful to develop recommendations for how to subsample datasets for best species tree inference. Here we apply a new measure of phylogenetic information content that estimates the reduction in tree space occupied by a posterior sample of inferred trees relative to a prior sample in order to assess the effects of gene tree parameters on species tree estimation. We find that, consistent with earlier studies, when data are informative, more data result in better species tree inference. However, when data are uninformative, subsampling a dataset to include only the most informative loci may produce a better species tree sample. We perform analyses on a variety of simulated and empirical datasets.

## Introduction

The current phylogenomics era is characterized by an abundance of genomic data along with the realization that topologically different phylogenies underlie different parts of the genome. One common source of incongruence among gene trees is incomplete lineage sorting (ILS), characterized by deep coalescences in gene trees (Maddison, 1997). The multispecies coalescent (MSC) model (Rannala and Yang, 2003) accommodates ILS, but Bayesian implementations are limited in the amount of data they can handle with a practical amount of computing resources. It is therefore useful to develop recommendations for how to subsample loci in phylogenomics data sets.

Measures to assess phylogenetic information content have been developed (Steel et al., 1993, 1995; Lyons-Weiler et al., 1996; Goldman, 1998; Massingham and Goldman, 2000; Shpak and Churchill, 2000; Xia et al., 2003; Xia, 2009; Geuten et al., 2007; Townsend, 2007; Shi et al., 2008; Fischer and Steel, 2009; Lemey et al., 2009; San Mauro et al., 2009; Tippery et al, 2012; Lewis et al., 2016; Townsend et al., 2012; Brown, 2014; Duchêne et al., 2022), but these measures have so far been applied to gene trees, not species trees (e.g. Prum et al., 2015). Additionally, these measures only take into account discrete tree topologies, not continuous parameters such as branch lengths. Because gene trees are often nuisance parameters required to estimate a species tree, it makes sense to assess how information about species trees is affected by various model parameters, providing insight into how to subsample loci for best species tree samples. Previous studies (Heled and Drummond, 2010; Corl and Ellegren, 2013) found that increasing the number of loci in a species tree analysis tends to improve resolution and accuracy of the species tree on both simulated and empirical data. However, these studies did not consider informativeness of the data in the analyses. Lanier et al. (2014) found high variation loci may produce a better species tree inference than low variation loci for a Bayesian framework. Lanier et al. (2014) used variation as a proxy for information content and did not estimate information content directly. This approach is problematic if high variation loci have low information content due to saturation (multiple substitutions per site) or low variation loci have high information content if sequences are long. Here, we estimate information content to assess whether including only the most informative loci in an analysis improves computational efficiency without sacrificing species tree accuracy.

We use a new method to calculate phylogenetic information content (*I*) that is described fully in Milkey and Lewis (2026). This method estimates the reduction in tree space occupied by a posterior sample of inferred trees relative to a prior sample. A large reduction in tree space corresponds to high information content.

Our method requires two samples of phylogenetic trees, one from the posterior and one from the prior. We estimate the mean tree (Miller, Owen, and Provan, 2015) from both the posterior and prior samples and calculate the geodesic distance (Owen and Provan, 2010) between the mean tree and each sampled tree in the BHV space defined by Billera et al. (2001). From these distances, we calculate the radius of a hypersphere that just includes the 95% set of sampled trees closest to the mean tree. Information content is defined as:

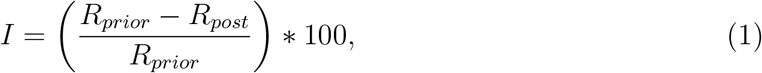

where *R*_*post*_ and *R*_*prior*_ are the radii for the posterior and prior samples, respectively.

Geodesic tree distances are functions of both branch lengths and topologies, so this measure of information content accounts for both tree characteristics. If the radius around the posterior mean tree is much smaller than the radius around the scaled prior mean tree, information content will approach 100%. If the radius around the posterior mean tree is equal to the radius around the scaled prior mean tree, information content will be 0%. This method may be applied to samples of gene trees or species trees.

We expect trees sampled under the prior to be much longer than trees sampled under the posterior because sequence data generally contain abundant information about branch lengths. To focus on topological information content and to prevent information about tree length from dominating, we scale the prior mean tree 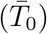 so it has the same total length as the posterior mean tree 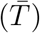. This scaling does not imply that only topological information is measured; correlations among edge lengths affect *I* even if all trees have the same overall tree length.

We use this measure of information content to assess the effect of different gene tree parameters on information about and accuracy of species trees using a variety of simulated datasets and one empirical dataset.

## Materials and Methods

We simulated all data sets and performed all Bayesian analyses using the program SMC, available at https://github.com/amilkey1/SMC, which samples from the MSC model using a sequential Monte Carlo approach (Milkey et al., 2025). Both the mean tree and tree distances were calculated in this paper using the open-source software op, available from https://github.com/plewis/op.

### Experiment 1: Sites per locus

At every point in a 10 by 10 grid, data were simulated under a Jukes-Cantor (Jukes and Cantor, 1969) model for 10 loci, 5 species, and 2 sampled individuals per species for a total of 100 simulated datasets. The 10 grid rows corresponded to evenly-spaced values of *T*, the species tree height, from 0.0 to 0.1. The 10 grid columns corresponded to evenly-spaced values of *θ/*2, from 0.0 to 0.1. *θ* was fixed across all species within a species tree. Smaller values of T and larger values of *θ/*2 yield greater expected deep coalescences, and thus those areas of the grid present a greater challenge to species tree methods.

We repeated the simulations 4 times for a total of 400 datasets, varying the number of sites per locus for each set of 100 datasets. For the first, second, and third sets, we simulated datasets with 10, 100, and 1000 sites per locus, respectively. For the fourth set of analyses, we conditioned species trees on the true gene trees (effectively an infinite number of sites per locus). We sampled species trees using the SMC program. See supplementary information for details about program settings. We calculated species tree information content (*I*) and inaccuracy. Here we measure inaccuracy as the average BHV distance between the sampled trees and the true tree. A smaller distance means the two trees are closer to one another in tree space, so a smaller BHV distance to the true tree indicates higher accuracy. We report inaccuracy as BHV distances between sampled trees and true tree but discuss these results in terms of accuracy rather than inaccuracy (i.e. lower BHV distance = higher accuracy). We expected species tree information and accuracy to increase with increasing per-locus sequence length.

### Experiment 2: Number of loci

We repeated the simulation conditions in experiment 1 but this time performed all analyses using the true gene trees as data. We simulated a different number of loci for each of 6 sets of 100 datasets (10, 20, 30, 40, 50, or 100 loci). We sampled species trees using the SMC program. See supplementary information for details about program settings. We calculated species tree information content (*I*) and inaccuracy (BHV distances from sampled trees to true tree). We expected species tree information and accuracy to increase with an increasing number of loci (given no gene tree estimation error).

### Experiment 3: Among-locus rate variation

We simulated 25 datasets under a Jukes-Cantor (Jukes and Cantor, 1969) model with fixed *T* = 0.1, fixed *θ* = 0.1, 100 loci, 5 species, 2 sampled individuals per species, and 1000 sites per locus. We divided loci into 9 groups, each with a different normalized relative rate, ranging from 0.0001 to 3.3. We sampled gene trees under the true model from both the prior and posterior using RevBayes v1.3.2 (Höhna et al., 2016) software. We calculated information content in each gene tree sample and conducted species tree analyses using only the most informative loci as determined by various cutoff points. To calculate gene tree information content, we applied our method to the gene tree samples. We expected locus informativeness to decrease at very low and very high rates. We calculated species tree information content (*I*) and inaccuracy (BHV average distances from sampled trees to true tree). We expected species tree information and accuracy to increase as more informative loci were used.

### Application: Empirical dataset

We assessed information content on an empirical dataset of teleost fishes (Near and Kim, 2021) with 22 species, 16 loci, 71 sampled taxa, and 12545 base pairs. We sampled gene trees under a GTR (Tavare and Miura, 1986) + I + G model from both the prior and posterior using RevBayes v.1.3.2 (Höhna et al., 2016) software. We calculated information content in each gene tree sample and conducted species tree analyses using only the most informative loci as determined by various cutoff points. Species tree analyses were performed in the SMC program using an HKY (Hasegawa et al., 1985) + I + G model (the SMC program does not currently implement a GTR model). Proportion of invariable sites, gamma shape parameter, and empirical base frequencies were estimated in PAUP* (Swofford, 2003). See supplementary information for details about priors and program settings. Because the maximum locus information content was roughly 51%, we chose cutoff points of 0% information (all 16 loci included in species tree analysis), 10% (only loci with *>*10% information included in species tree analysis), 20%, 30%, 40%, and 50%. We did not use a cutoff above 50% because this would have removed all loci from the analysis. For computational efficiency, we estimated a prior mean tree for the first locus and used this as the prior mean tree for all loci. We calculated species tree information content (*I*) but because this is an empirical dataset we were not able to calculate species tree accuracy.

## Results

### Experiment 1: Amount of sequence data

Information about species trees and accuracy of species trees generally increased with increasing per-locus sequence length (Fig. 1), with the most dramatic improvement from 10 sites to 100 sites. Information content for the 10-, 100-, 1000-, and infinite-site analyses was on average 60.6%, 76.7%, 86.5%, and 93.3%, respectively. Inaccuracy for the analyses was on average 0.056, 0.034, 0.022, and 0.007, respectively. Information and accuracy tended to decrease with an increase in the proportion of deep coalescences (Fig. 1). Even with an infinite number of sites per locus, perfect accuracy is not expected with only 10 loci.

**Figure 1.**
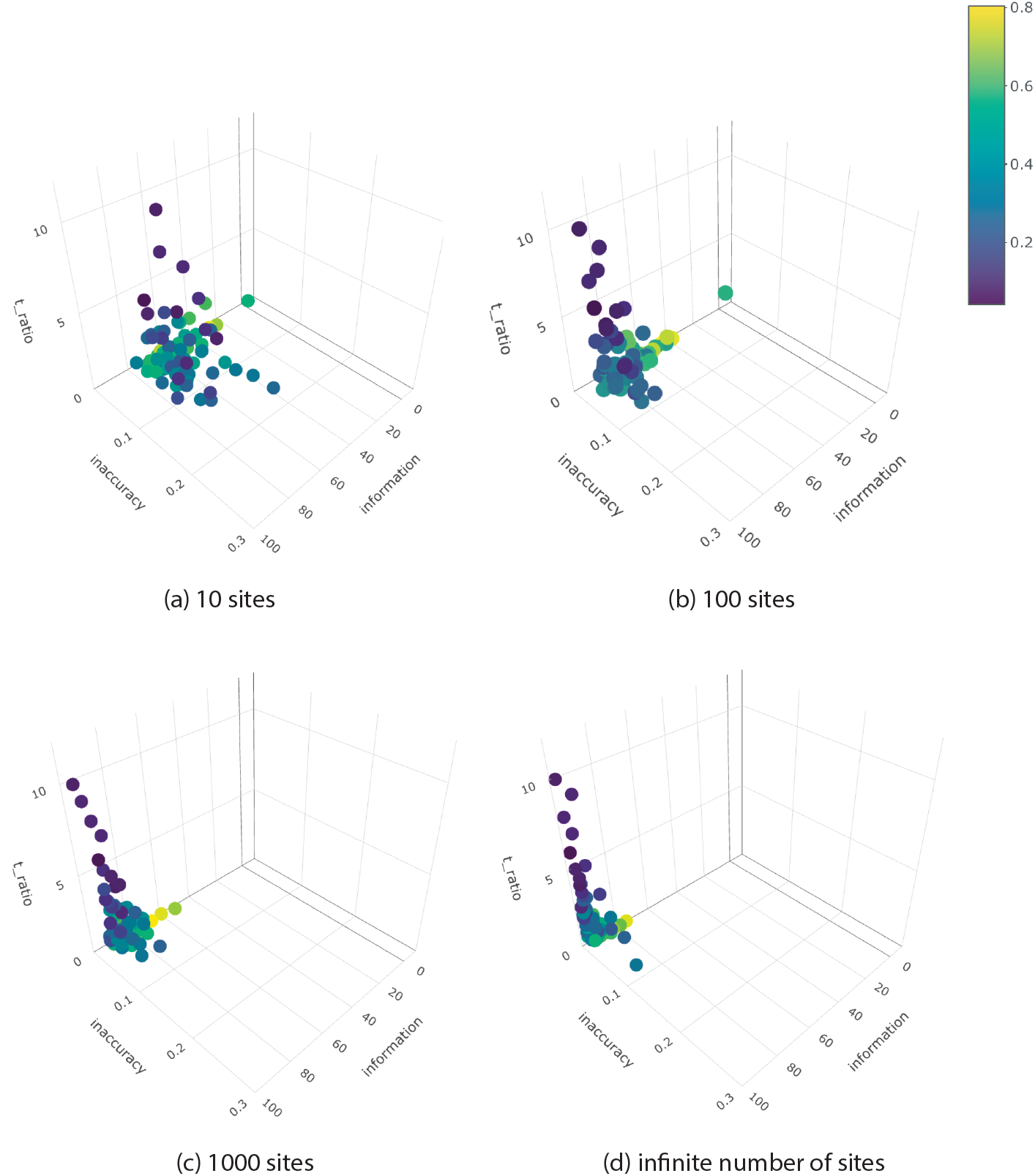
Experiment 1 results (10 loci, 5 species, 2 sequences sampled for each species). Analyses performed with varying amounts of sequence data simulated. a) 10 sites per locus. b) 100 sites per locus. c) 1000 sites per locus. d) Infinite number of sites (species trees conditioned on true gene trees). Colors indicate proportion of all coalescences that are deep across all gene trees. Axes represent information content (topology and branch lengths), inaccuracy (BHV distances), and ratio of species tree height to average coalescence time.

### Experiment 2: Number of loci

Information about species trees and accuracy of species trees generally increased with an increasing number of loci (Fig. 2). Information content for the 10-, 20-, 30-, 40-, 50-, and 100-locus analyses was on average 96.6%, 98.5%, 99.1%, 99.4%, 99.5%, and 99.8%, respectively. Inaccuracy for the analyses was on average 0.0064, 0.0038, 0.0028, 0.0025, 0.0024, and 0.0020, respectively.

**Figure 2.**
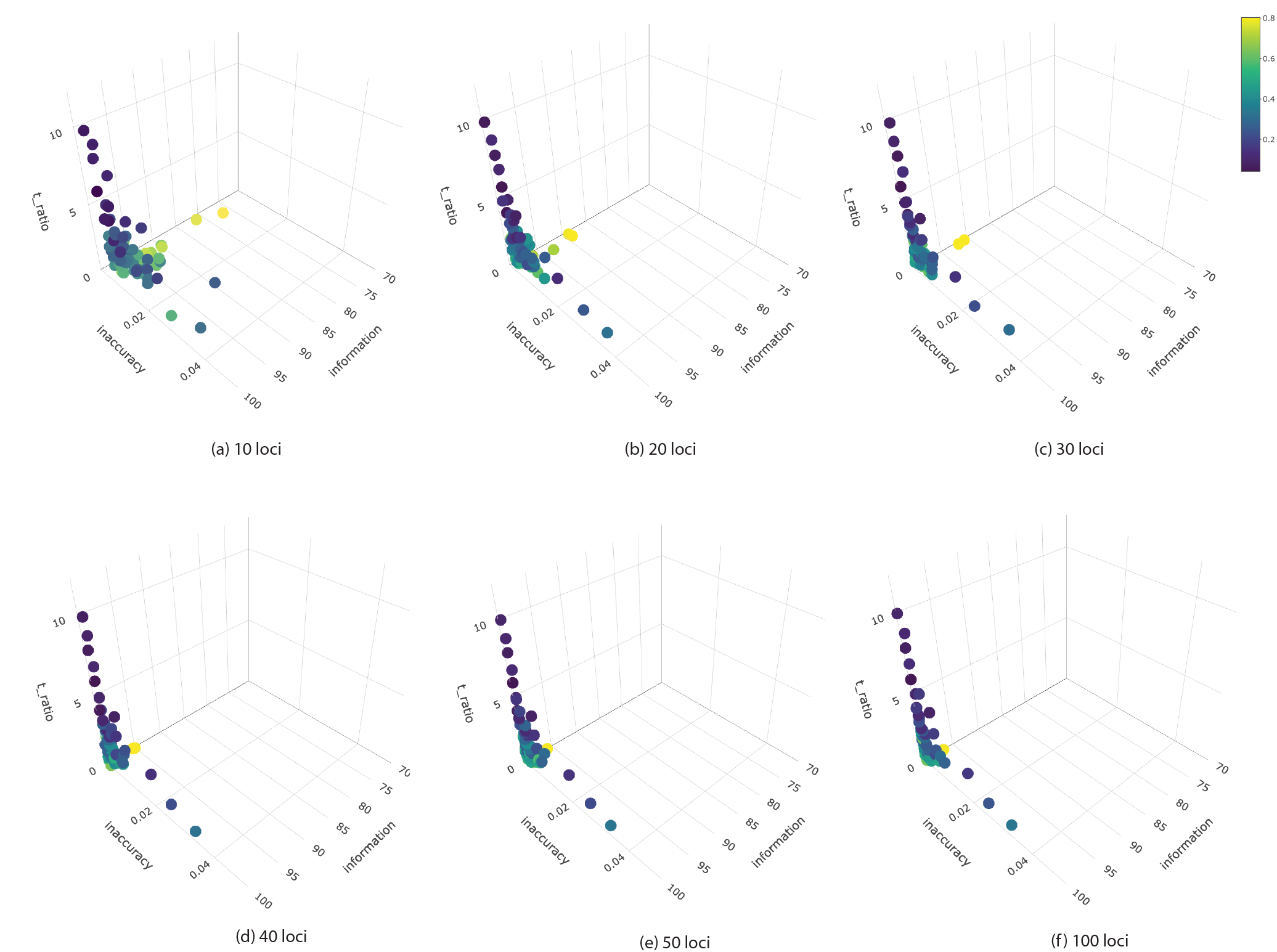
Experiment 2 results (5 species, 2 sequences sampled each species, infinite number of sites each locus). Analyses performed with varying numbers of loci simulated. a) 10 loci. b) 20 loci. c) 30 loci. d) 40 loci. e) 50 loci. f) 100 loci. For all analyses, species trees were conditioned on true gene trees. Colors indicate proportion of all coalescences that are deep across all gene trees. Axes represent information content (topology and branch lengths), inaccuracy (BHV distances), and ratio of species tree height to average coalescence time.

Information content and accuracy were both lowest for the 1-locus datasets (Fig. 2). Information and accuracy tended to decrease with an increase in the proportion of deep coalescences (Fig. 2). While information content and accuracy were highest for the 100-locus datasets, there were only very marginal improvements over these measures on the 30-, 40-, and 50-locus datasets.

### Experiment 3: Among locus rate variation

Locus information content peaked at 92.8% at the relative rate of 2.0 but was very high for all relative rates between 1.0 and 3.3 (Fig. 3a). Locus information content was very low at low rates of evolution (just 0.7% information at relative rate of 0.0001, 8.1% information at relative rate of 0.001, 42.8% information at relative rate of 0.01, and 78% information at relative rate of 0.1). High rate sites may have lower information due to the noise of substitutional saturation, though we did not choose relative rates high enough to see a reduction in information at high rates. Low rate sites have lower information due to a paucity of variable sites. This experiment illustrates that high rate loci are more informative than low rate loci. That is, it is better to have more variable sites than to simply have no variation at all.

**Figure 3.**
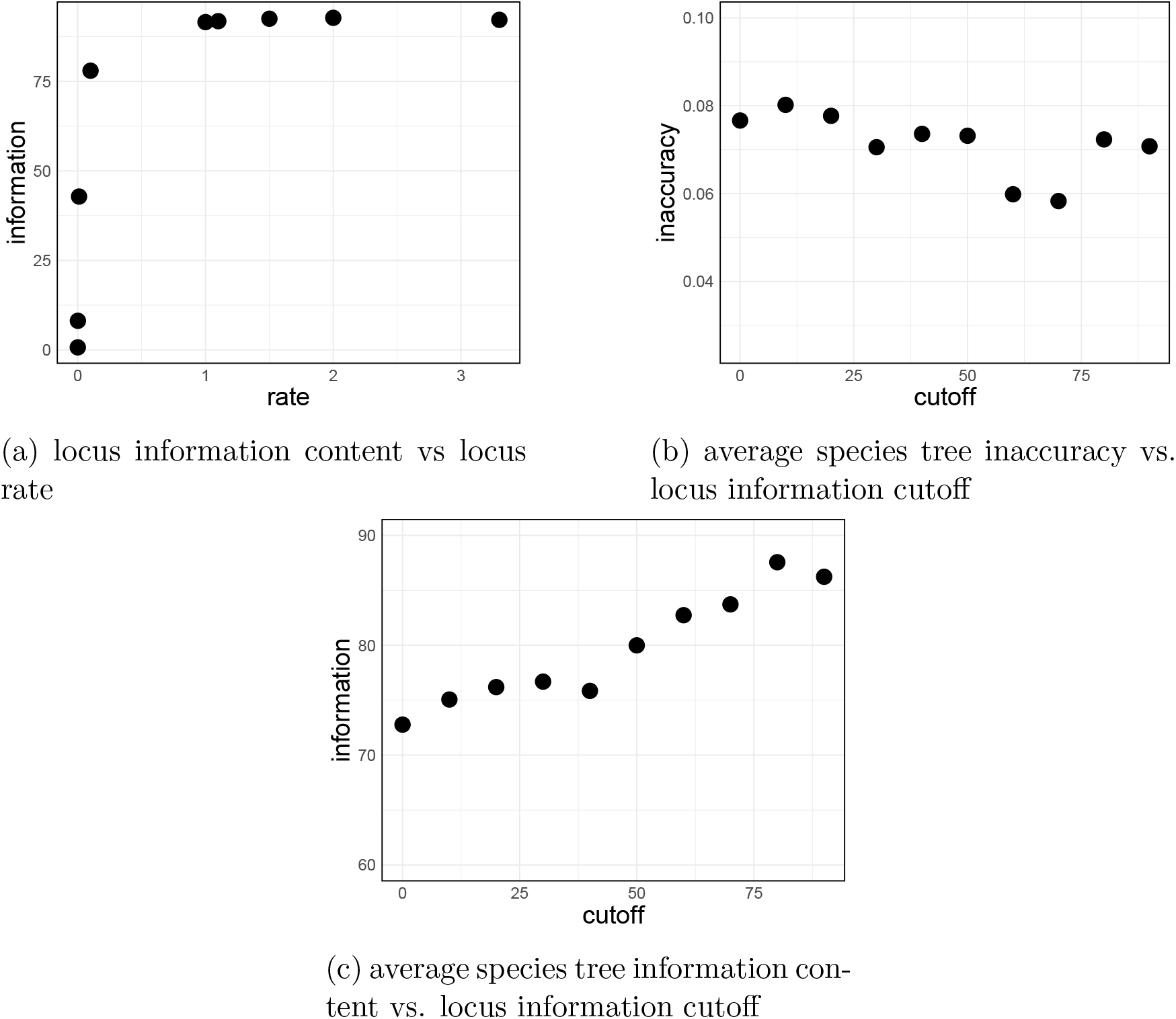
Experiment 3 results (*T* = 0.1, *θ* = 0.1, 100 loci, 5 species, and 2 sampled sequences each species). a) Locus information content vs locus rate. b) Average sampled species tree inaccuracy vs locus information cutoff. c) Average information about species trees vs locus information cutoff.

Species tree accuracy was highest (BHV distance 0.058) at a locus information cutoff of 70% (i.e. a locus must have at least 70% information to be included), despite this cutoff leaving fewer loci available for analyses than cutoff values 0% through 60% (Fig. 3b). Species tree accuracy was lowest (BHV distance 0.080) when the locus-specific cutoff was 10% (all but the least informative loci included in the analysis). While there was a slight decrease in species tree accuracy at the 80% and 90% locus cutoffs, in general, species tree accuracy increased with an increase in the locus-specific information cutoff. Information about species trees was highest (87.5%) when the locus-specific information cutoff was high (80%) and lowest (72.8%) when the locus-specific information cutoff was 0% (all loci included in the analysis). Information about species trees increased consistently with an increase in locus-specific information cutoff with the exception of slight drops at the 50% and 90% cutoffs.

### Empirical dataset

Average information content across all 16 loci was 31.69%. The lowest locus information content was 6.83% (locus 12), and the highest locus information content was 51.4% (locus 2). Our cutoff values therefore ranged from 0% information (no loci removed) to 50% information (15 loci removed) (Fig. 4b).

**Figure 4.**
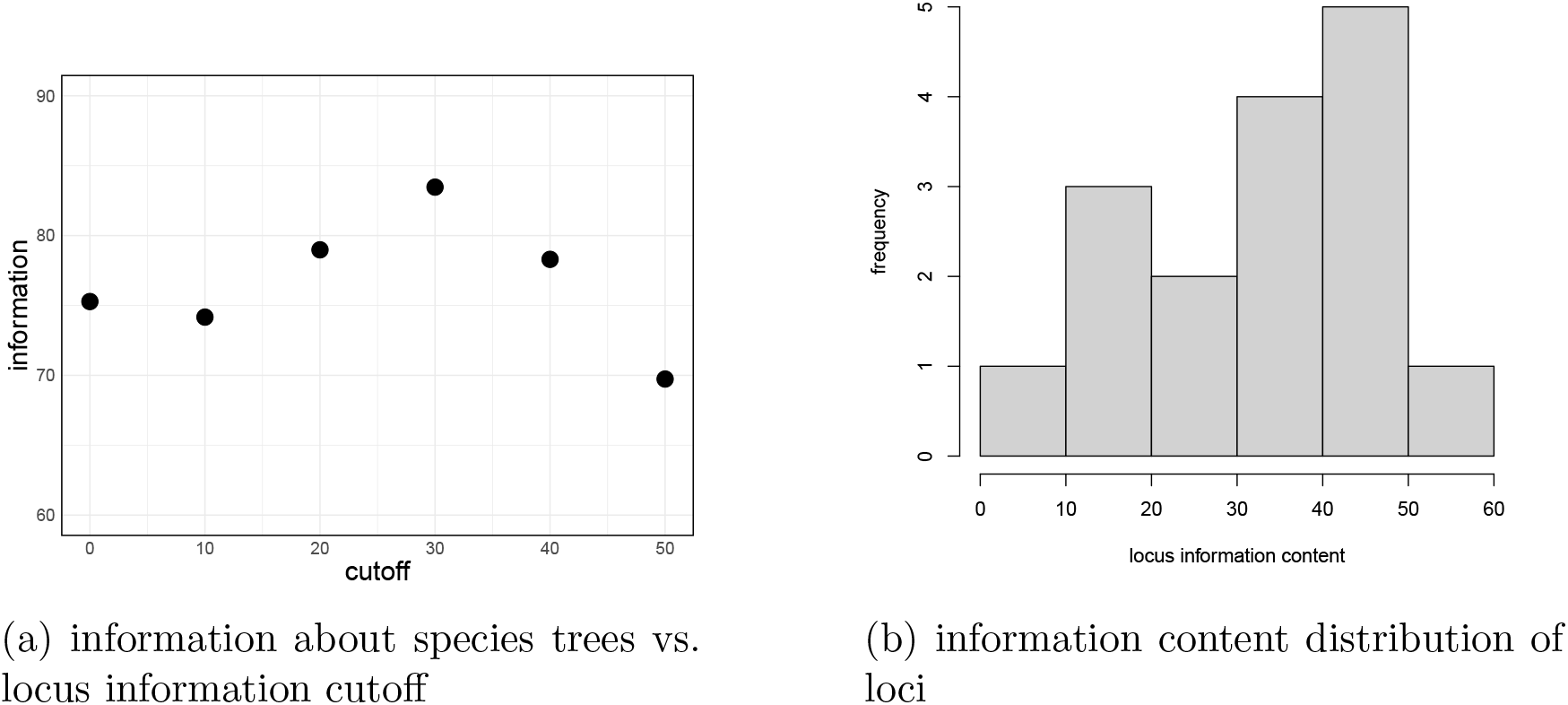
Empirical results. a) Information about species trees vs locus information cutoff. b) Information content distribution of 16 loci.

Species tree information content decreased slightly from the 0% locus information cutoff to the 10% locus information cutoff and then increased through the 30% information cutoff.

Species tree information content dropped to its lowest at the 50% information cutoff (Fig. 4a). The 50% cutoff analyses only included 1 locus, and the 40% information cutoff analyses only included 6 loci, compared to 10 or more loci for the other analyses.

## Discussion

How much data is needed to accurately resolve a phylogeny is a common question in phylogenetics. Increasing the number of loci from a few loci to a large number of loci may not improve a species tree in practice or may make the inference worse. Furthermore, large datasets run into computational limits, which has made downsampling a common strategy for Bayesian phylogenetic analyses. This raises questions about how much information data actually contain about species trees and how to best subsample loci for an informative species tree sample, which we addressed here.

Experiments 1 and 2 demonstrate that, when data are informative, adding more data to an analysis (more sequences or more loci) results in a better species tree inference. However, experiment 2 showed there may be little benefit to continuing to add more data to an analysis when a good species tree sample has already been achieved. While experiment 2 was conducted with the best quality data possible (true gene trees as data), which is not realistic for empirical data, the very minor increase in species tree accuracy from 30 loci to 100 loci demonstrated there was no practical value to adding more data to this analysis. It is worth keeping in mind that, while the multispecies coalescent model is statistically consistent (Allman et al., 2011; Mirarab, Nakhleh, and Warnow, 2021) and should therefore see increased species tree accuracy with more data if no parts of the model are violated, very slight improvements in species tree inference with more data may not be worth the extra computational resources needed to run the model with the extra data. We therefore suggest focusing on using the most informative loci, as determined by a comparison of posterior and prior gene tree samples.

Experiment 3 demonstrates that, when loci are uninformative, including more loci may actually reduce the accuracy of the species tree. We found that slow-evolving loci in particular had lowest levels of information due to few substitutions, and the resulting species trees were more accurate when these loci were removed from the analysis. This contradicts existing ideas that more loci always improve the species tree inference (Heled and Drummond, 2010; Corl and Ellegren, 2013). We also found that information about and accuracy of species trees generally increased as we increased the gene tree information cutoff and jumped to its highest levels when the least informative loci were excluded. These results suggest it may be important to consider locus information content when deciding which loci to include in a species tree analysis.

While our empirical analysis was less decisive than the simulation studies, it similarly showed that removing the least informative loci tended to increase information about the species tree. This analysis also demonstrated the pitfalls of removing too many loci. We found a slight decrease in species tree information when we only included the 6 most informative loci (gene tree information cutoff greater than 40%) as opposed to the 10 most informative loci (gene tree information cutoff greater than 30%), which indicates 6 loci may not be enough for a good species tree inference in this case. We found a large decrease in species tree information when we only included the single most informative locus (gene tree information cutoff greater than 50%); a single gene tree is rarely sufficient for good species tree inference.

The inference program used for analyses (Milkey et al., 2025) involves estimating gene trees in order to estimate the species tree, as do many other species tree programs, such as StarBEAST3 (Douglas et al., 2014) and ASTRAL (Mirarab and Warnow, 2015). We did not assess information content under methods that integrate out gene trees, such as SNAPP (Bryant et al., 2012) and SVDQuartets (Chifman and Kubatko, 2014). For particularly low-information loci (e.g. slow-evolving or short loci), methods that bypass gene tree estimation by effectively accounting for all possible gene trees may be useful as they are better able to take advantage of all the information in the data.

### Recommendations

For species tree methods that involve gene tree estimation (e.g. Milkey et al., 2025; Douglas et al., 2014; Mirarab and Warnow, 2015), we recommend assessing locus information content and removing loci with very low information.

This appears to contradict the assertion in Lanier et al. (2014) that a biological justification should be used as a guide to excluding loci, not a lack of information; however, Lanier et al. (2014) did not measure information content, instead using substitution rate as a proxy. In contrast, we directly measured information content and used a diversity of simulations to demonstrate that species tree inference generally improves with the exclusion of low-information loci.

In addition, if locus sequence length is consistently very short, it may be difficult to estimate gene trees accurately (as we showed in experiment 1). In this case, we recommend considering a site-based method (e.g. Chifman and Kubatko, 2014) that does not estimate gene trees (Molloy and Warnow, 2018). We urge readers to carefully consider informativeness of data when conducting an analysis and to not assume that more data will necessarily result in a better species tree inference.

## Acknowledgements

We would like to thank the Associate Editor and two anonymous reviewers for their constructive comments and suggestions. We would like to thank Elizabeth Jockusch and Jill Wegrzyn for their constructive comments.

## Supplementary Material

Supplementary material can be found at DOI 10.5281/zenodo.19359292.

## References

Allman, S. E, H. J. H. Degnan, and J. A. Rhodes. 2011. Identifying the rooted species tree from the distribution of unrooted gene trees under the coalescent. Journal of Mathematical Biology 62(6):833–862. 10.1007/s00285-010-0355-7

Billera, L. J., S. P. Holmes, and K. Vogtmann. 2001. Geometry of the space of phylogenetic trees. Advances in Applied Mathematics 27:733–767. 10.1006/aama.2001.0759

Brown, J. M. 2014. Detection of implausible phylogenetic inferences using posterior predictive assessment of model fit. Systematic Biology 63:334–348. 10.1093/sysbio/syu002

Bryant, D., R. Bouckaert, J. Felsenstein, N. A. Rosenberg, and A. RoyChoudhury. 2012. Inferring species trees directly from biallelic genetic markers: bypassing gene trees in a full coalescent analysis. Molecular Biology and Evolution 29:1917–1932. 10.1093/molbev/mss086

Chifman, J., and L. Kubatko. 2014. Quartet inference from SNP data under the coalescent model. Bioinformatics 30:3317–3324. 10.1093/bioinformatics/btu530

Corl, A., and H. Ellegren. 2013. Sampling strategies for species trees: the effects on phylogenetic inference of the number of genes, number of individuals, and whether loci are mitochondrial, sex-linked, or autosomal. Molecular Phylogenetics and Evolution 67:358– 366. 10.1016/j.ympev.2013.02.002

Douglas, J., C.L. Jiménez-Silva, and R. Bouckaert. 2022. StarBeast3: adaptive parallelized Bayesian inference under the multispecies coalescent. Systematic Biology 71:901– 916. 10.1093/sysbio/syac010

Duchêne, D. A., N. Mather, C. Van Der Wal, and S. Y. W. Ho. 2022. Excluding loci with substitution saturation improves inferences from phylogenomic data. Systematic Biology 71:676-–689. 10.1093/sysbio/syab075

Fischer, M., and M. Steel. 2009. Sequence length bounds for resolving a deep phylogenetic divergence. Journal of Theoretical Biology 256:247–252. 10.1016/j.jtbi.2008.09.031

Geuten, K., T. Massingham, P. Darius, E. Smets, and N. Goldman. 2007. Experimental design criteria in phylogenetics: where to add taxa. Systematic Biology 56:609–622. 10.1080/10635150701499563

Goldman, N. 1998. Phylogenetic information and experimental design in molecular systematics. Proceedings of the Royal Society of London Series B 265:1779–1786. 10.1098/rspb.1998.0502

Jukes, T. H., and C. R. Cantor. 1969. Evolution of protein molecules. In Munro, H. N., editor. Mammalian Protein Metabolism. New York: Academic Press. 21–132. https://evoluscope.fr/phylographe/biblio/JukesCantor1969.pdf

Hasegawa, M., H. Kishino, and T. Yano. 1985. Dating of the human-ape splitting by a molecular clock of mitochondrial DNA. Journal of Molecular Evolution 2(2):160–174. 10.1007/BF02101694

Heled, J., and A. J. Drummond. 2010. Bayesian inference of species trees from multilocus data. Molecular Biology and Evolution 27:570–580. 10.1093/molbev/msp274

Höhna, S., M.J. Landis., T. A. Heath, B. Boussau, and F. Ronquist. 2016. RevBayes: Bayesian phylogenetic inference using graphical models and an interactive model-specification language. Systematic Biology 65:726–736. 10.1093/sysbio/syw021

Lanier, H. C., H. Huang, and L. L. Knowles. 2014. How low can you go? The effects of mutation rate on the accuracy of species-tree estimation. Molecular Phylogenetics and Evolution 70:112–119. 10.1016/j.ympev.2013.09.006

Lemey, P., A. Rambout, A. J. Drummond, and M. A. Suchard. 2009. Bayesian phylogeography finds its roots. PLoS Computational Biology 5:e1000520. 10.1371/journal.pcbi.1000520

Lewis, P. O., M. H. Chen, L. Kuo, L. A. Lewis, K. Fučíková, S. Neupane, Y. B. Wang, and D. Shi. 2016. Estimating Bayesian phylogenetic information content. Systematic Biology 65:1009–1023. 10.1093/sysbio/syw042

Lyons-Weiler, J., G. A. Hoelzer, and R. J. Tusch. 1996. elative apparent synapomorphy analysis (RASA). I: The statistical measurement of phylogenetic signal. Molecular Biology and Evolution 13:749–757. 10.1093/oxfordjournals.molbev.a025635

Maddison, W. 1997. Gene trees in species trees. Systematic Biology 46:523–536. 10.1093/sysbio/46.3.523

Massingham, T., and N. Goldman. 2000. EDIBLE: experimental design and information calculations in phylogenetics. Bioinformatics 16:294–295. 10.1093/bioinformatics/16.3.294

Miller, E., M. Owen, and J. S. Provan 2015. Polyhedral computational geometry for averaging metric phylogenetic trees. Advances in Applied Mathematics 68:51–91. 10.1016/j.aam.2015.04.002

Milkey, A., M. H. Chen, Y. B. Wang, A. Li, and P. O. Lewis. 2025. The sequential multispecies coalescent. bioRxiv. https://www.biorxiv.org/content/10.1101/2025.01.31.635964v1

Milkey, A., and P. O. Lewis. 2026. Estimating Bayesian phylogenetic information content using geodesic distances. bioRxiv. https://www.biorxiv.org/content/10.64898/2026.03.31.715656v1

Mirarab, S., and T. Warnow. 2015. ASTRAL-II: Coalescent-based species tree estimation with many hundreds of taxa and thousands of genes. Bioinformatics 31:i44–i52. 10.1093/bioinformatics/btv234

Mirarab, S., L. Nakhelh, and T. Warnow. Multispecies coalescent: theory and application in phylogenetics. Annual Review of Ecology, Evolution, and Systematics 52(1):247–268. 10.1146/annurev-ecolsys-012121-095340

Molloy, K., and T. Warnow. 2018. To include or not to include: the impact of gene filtering on species tree estimation methods. Systematic Biology 67:285–303. 10.1093/sysbio/syx077

Near, T. J., and D. Kim. 2021. Comparison of phylogenetic trees. Phylogeny and time scale of diversification in the fossil-rich sunfishes and black basses (Teleostei: Percomorpha: Centrarchidae). Molecular Phylogenetics and Evolution 161:107156. 10.1016/j.ympev.2021.107156

Owen, M., and J. S. Provan. 2010. A fast algorithm for computing geodesic distances in tree spaces. IEEE/ACM Transactions on Computational Biology and Bioinformatics 8:2–13. 10.1109/TCBB.2010.3

Prum, R. O., J. S. Berv, A. Dornburg, D. J. Field, J. P. Townsend, E. M. Lemmon, and A. Lemmon. R. 2015. A comprehensive phylogeny of birds (Aves) using targeted next-generation DNA sequencing. Nature 526:569–573. 10.1038/nature15697

Rannala, B., and Z. Yang. 2003. Bayes estimation of species divergence times and ancestral population sizes using DNA sequences from multiple loci. Genetics 164:1645–1656. 10.1093/genetics/164.4.1645

San Mauro, D., D. J. Gower, T. Massingham, M. Wilkinson, R. Zardoya, J. A. Cotton. 2009. Experimental Design in Caecilian Systematics: Phylogenetic Information of Mitochondrial Genomes and Nuclear rag1. Systematic Biology 58:425–438. 10.1093/sysbio/syp043

Steel, M., P. J. Lockhart, and D. Penny. 1993. A frequency-dependent significance test for parsimony. Molecular Phylogenetics and Evolution 4:64–71. 10.1006/mpev.1995.1006

Steel, M., P. J. Lockhart, and D. Penny. 1995. Confidence in evolutionary trees from biological sequence data. Nature 364:440–442. 10.1038/364440a0

Shi, X., H. Gu, and C. Field. 2008. Pattern Classification of Phylogeny Signals. Statistical Applications in Genetics and Molecular Biology 7:1. 10.2202/1544-6115.1289

Shpak, M., and G. A. Churchill. 2008. The information content of a character under a Markov model of evolution. Molecular Phylogenetics and Evolution 17:231–243. 10.1006/mpev.2000.0846

Swofford D. L. 2003. PAUP*. Phylogenetic Analysis Using Parsimony (*and Other Methods. Version 4). Sinauer Associates. http://paup.phylosolutions.com

Tavare S. and R. M. Miura. 1986. PAUP*. Some probabilistic and statistical problems in the analysis of DNA sequences. Lectures on Mathematics in the Life Sciences. 17:57–86.

Tippery, N. P., K. Fučíková, P. O. Lewis, and L. A. Lewis. 2012. Probing the monophyly of the Sphaeropleales (Chlorophyceae) using data from five genes. Journal of Phycology 48:1482–1493. 10.1111/jpy.12003

Townsend, J.P. 2007. Profiling phylogenetic informativeness. Systematic Biology 56:222–231. 10.1080/10635150701311362

Townsend, J. P., Z. Su, and Y. I. Tekle. 2012. Phylogenetic signal and noise: predicting the power of a data set to resolve phylogeny. Systematic Biology 61:835–849. 10.1093/sysbio/sys036

Xia, X., Z. Xie, M. Salemi, L. Chen, and Y. Wang. 2003. An index of substitution saturation and its application. Molecular Phylogenetics and Evolution 26:1–7. 10.1016/S1055-7903(02)00326-3

Xia, X. 2009. Assessing substitution saturation with DAMBE. Pages 613–629 in: Lemey, P., M. Salemi, and A.-M. Vandamme. (eds.). The phylogenetic handbook: a practical approach to phylogenetic analsysis and hypothesis testing. Cambridge University Press. 2nd edition. 10.1017/CBO9780511819049

